# TomoSegNet: Augmented membrane segmentation for cryo-electron tomography by simulating the cellular context

**DOI:** 10.64898/2026.01.15.699326

**Authors:** Rachida Seghiri, Juan Diego Gallego Nicolás, Robert Brandt, Miguel A. Meroño, Pierre Lefevre, Noushin Hajarolasvadi, Daniel Baum, Jessica Heebner, Harold Phelippeau, Pascal Doux, Antonio Martinez-Sanchez

**Affiliations:** Advanced Technology, Thermo Fisher Scientific, 39 Rue d’Armagnac, Bordeaux, 33800, France; Department of Information and Communications Engineering, Universidad de Murcia, Campus de Espinardo, Murcia, 30100, Spain; Department of Mathematics, Universidad de Murcia, Campus de Espinardo, Murcia, 30100, Spain; Department of Visual and Data-Centric Computing, Zuse Institute Berlin, Takustr. 7, Berlin, 14195, Germany

**Author notes:** Contributing authors. These authors contributed equally to this work (Rachida Seghiri, Juan Diego Gallego Nicolás).

**Keywords:** Cryo-Electron Tomography, Segmentation, Cellular Membrane, Image Processing, Synthetic data

## Abstract

Membrane segmentation is an essential task in the workflow for processing cryo-electron tomography data. Recently, machine learning algorithms have successfully been adopted to perform membrane segmentation. However, the performance of these approaches is limited by the training dataset, as models are trained from manual annotations, thereby hindering the models’ generalization and preventing the recovery of membranes that have vanished due to distortions. Here, we address these limitations by generating training data with a simulator. To provide a representative and realistic dataset, we have extended the current state-of-the-art in simulators for cryo-electron tomography by incorporating a biophysical model for membranes. We demonstrate that our machine learning model, trained solely from synthetic data, and thanks to the physical knowledge learned from the simulator, outperforms the current state-of-the-art for membrane segmentation in a diverse set of experimental data. This performance is particularly noteworthy in terms of recovering membranes lost due to imaging distortions.

## 1 Introduction

Biological membranes are the primary structural component of the cell, defining the cell boundary and its internal compartments. In addition, they are decorated with numerous membrane-bound molecular complexes that play a key role in many cellular processes. Cryo-Electron Tomography (cryo-ET) can reconstruct 3D tomograms at nanometer resolution from 2D micrographs, preserving the structure of the membranes and their molecular constituents [1]. However, the elevated levels of noise and imaging defects, together with the high density of heterogeneous structures within the cell and the complexity of the membrane’s geometries, make membrane visualization and analysis cumbersome. A specific distortion in electron tomography, named missing wedge, is particularly harmful to membranes. Since the sample is tilted in a limited range of angles, typically [− 60, 60]^*◦*^ or less, during micrograph acquisition, membranes with normals beyond these tilting angles tend to vanish [2].

Computer methods for membrane segmentation are required for the visualization and the quantitative analysis of the membrane structure [3, 4] as well as the associated proteins [5–7]. First attempts for membrane segmentation in cryo-ET were based on template matching algorithms [8, 9], but their computational cost and performance limit their application. TomoSegMemTV [10] drastically reduces the computational cost by applying steerable filters and increases the robustness by using the Tensor Voting algorithm [11, 12]. TomoSegMemTV was the first computational method robust enough to process cryo-electron tomograms, being the baseline for current developments. TomoSegMemTV is, in essence, a detector for surface-like structures, but it has two limitations. First, its output requires manual curation because non-membranous surface-like structures are also segmented. Second, vanishing membranes due to the missing wedge are not recovered.

Deep learning has emerged as the natural solution to overcome the limitations of analyzing cryo-electron tomograms. 3D-UNet [13], based on convolutional neural networks (CNN), is the most widely used architecture for macromolecular detection in 3D cryo-electron tomograms [14–16]. Machine learning methods require annotated data for training a model. MemBrain-Seg [17], also based on 3D-UNet, has been trained mainly with an extensive annotated dataset providing more continuous segmentation than non-machine learning approaches. TARDIS [18] uses a vision-transformer trained with diverse datasets and allows instance segmentation. Nevertheless, current deep learning approaches have two limitations. First, their performance depends on the biological sample; despite the efforts to include tomograms from diverse sources, every biological sample has particularities that may not be observed during the training. Second, manual annotations are inherently incomplete and may contain errors; for instance, the missing wedge artifact can render membranes invisible, preventing human annotators from labeling them. Consequently, here we propose to use synthetic data in order to obtain a perfectly characterized training set with a high diversity of structural features.

The past decade of machine learning research has consistently demonstrated that the improvements in model performance are driven by the scale, diversity, and quality of training data rather than architectural innovations alone. As an example, CNN architectures were proposed some decades ago [19], but their potential was demonstrated much later thanks to the ImageNet dataset [20], which also contributed to their subsequent development. By contrast, in the domain of cryo-ET membrane segmentation, the scarcity of accurately annotated data represents a fundamental bottleneck for training robust deep learning models. Traditional approaches rely heavily on manual annotation, which is not only time-consuming and labor-intensive but also prone to human error and inconsistencies, considering the low SNR and complex 3D structures present in cryo-ET. Moreover, the missing wedge artifact inherent in cryo-ET data means that even expert annotations may fail to capture the complete membrane structure, leading to incomplete ground truth labels that can negatively impact model performance. To overcome these limitations, we propose a data-centric AI framework for the design of synthetic training data using PolNet [21], enabling more accurate, generalizable, and artifact-aware deep learning models for segmentation of cryo-ET data. Before this article, PolNet could only generate a few simple geometrical shapes. However, these geometrical models were still too simple compared to the complexity of membranous organelles such as the endoplasmic reticulum.

Here, we present TomoSegNet, which overcomes the current limitations of deep learning methods for membrane segmentation: generalization and the recovery of membranes faded by the missing wedge. The package is available upon request via email, either as a standalone Python package with a command-line interface (CLI) or through a graphical user interface implemented as an Amira plugin with a time-limited free license. We also present an extension of PolNet (https://github.com/anmartinezs/polnet) simulator, which generates considerably more realistic membranes, a key factor in improving TomoSegNet performance. In addition, we carry out a study to determine the key elements of the synthetic data. Finally, we validate our method with an extensive set of experimental data and compare it visually and quantitatively with the state-of-the-art methods like MemBrain-Seg and TARDIS.

## 2 Results

### 2.1 A synthetic-data-centric AI framework

Broadly speaking, deep learning workflows are either model-centric or data-centric. Model-centric strategies iterate architectures and training schemes against fixed benchmarks, whereas data-centric strategies prioritize the quality, diversity, and representativeness of datasets. Currently, most deep learning methods for bioimaging align with model-centric strategies [22]. For instance, a recent benchmark dataset was introduced to provide a standardized dataset for the development and comparison of particle picking models [23]. In practice, data-centric approaches are more suitable for bioimaging [22], especially in settings lacking robust benchmarks such as membrane segmentation in cryo-ET. Moreover, data-centric approaches are more resilient to the rapid advances of cryo-ET in terms of resolution, automation, sample preparation, and reconstruction, which can quickly outdate fixed-benchmark models.

Motivated by these considerations, we introduce a data-centric AI framework for cryo-ET membrane segmentation that iteratively refines a synthetic-only training set while evaluating on experimental (real) data (Fig. 1). Departing from a conventional train/validation/test split, we assign distinct roles to three datasets: a synthetic-only training set; an experimental validation set, annotated by an expert and used for error analysis; and an experimental test set held back for a single, leakage-free assessment of generalizability [24]. At the core of the framework is an iterative development workflow: **G**enerate/refine synthetic data, **T**rain a model, **A**ssess systematic miss-predictions to define the required training data refinements. The loop repeats, with a human-in-the-loop, until the performance meets the target. Biology experts help surfacing failure modes and hard cases; these findings inform hypotheses about missing variability or physics in the synthetic generator. PolNet is then extended or re-tuned (e.g., membrane morphology, imaging noise, reconstruction, missing-wedge parameters), and a more representative training set is generated. We document four development cycles (V1–4) for membrane segmentation, corresponding to four versions of the synthetic training sets, with performance on the validation set summarized in Fig. 2.A. Coupling training data synthesis to empirical model performance progressively aligns the synthetic distribution with the complexity and variability of experimental tomograms. For example, after the second development cycle (Fig. 1), we implemented a more realistic membrane module in PolNet (Sec. 2.4), which was pivotal in enabling TomoSegNet to outperform models trained on manual annotations. A detailed description of the synthetic datasets V1-4 is available in Methods 4.2.

**Fig. 1.**
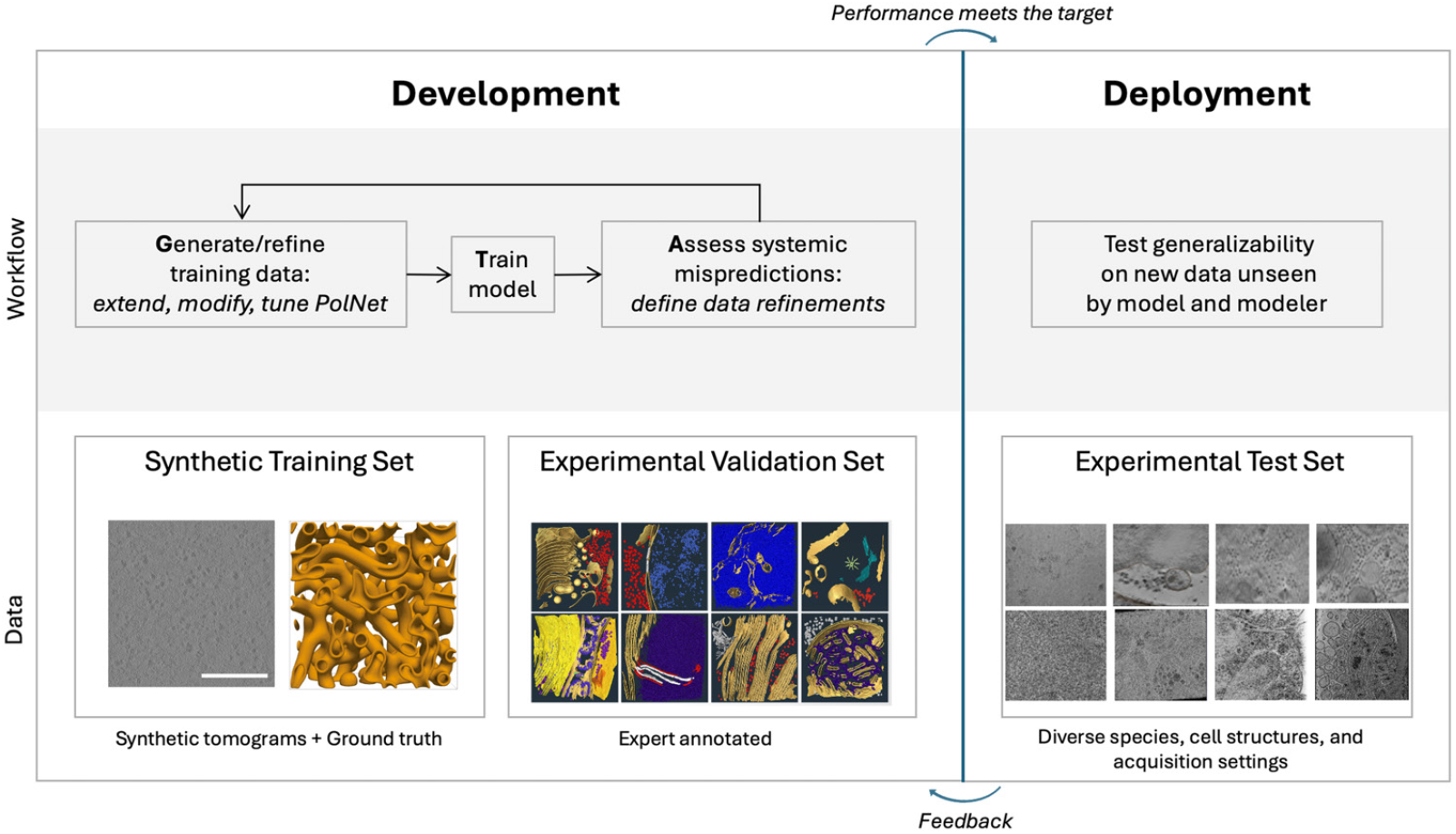
A synthetic-data-driven AI framework for cryo-ET. The framework couples data synthesis to empirical model performance. Deliverables include a generalizable model, high-quality synthetic cryo-ET tomograms, and expanded PolNet simulator capabilities. The relative size of workflow steps indicates the effort invested.

**Fig. 2.**
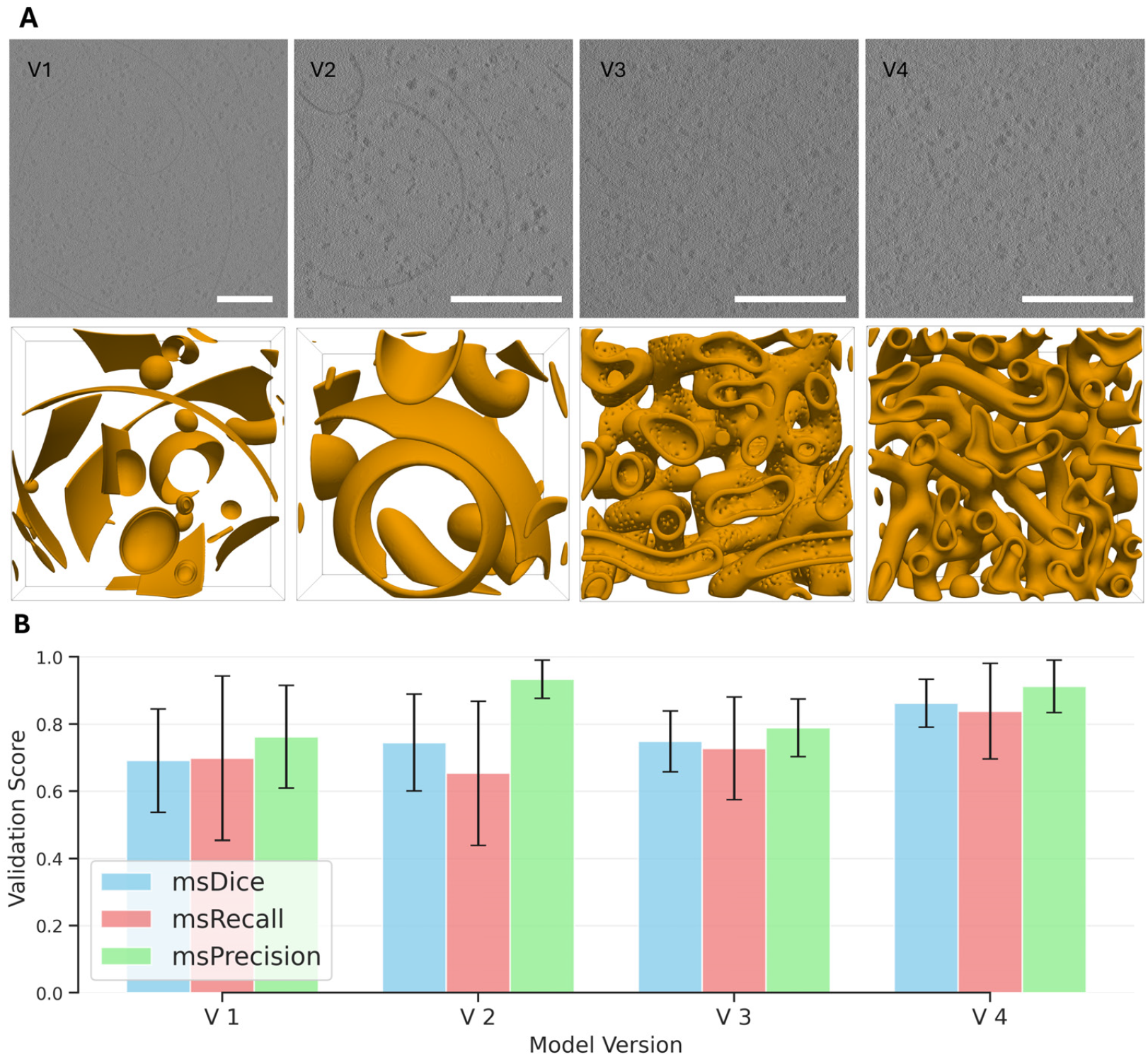
Evolution of the synthetic datasets. (A) Exemplary 2D slices (top row) and a 3D views (bottom row) of the membrane ground truth of the four synthetic dataset versions used for training. V1-V4 from left to right. Scale bars: 200 nm. (B) Evaluation of the successive model versions on the experimental validation set, described in Methods 4.1. We report msDice, msRecall, and msPrecision scores (± 1 std).

For training TomoSegNet, we adopt nnU-Net [25], a well-established baseline in medical image segmentation [26] that accelerates iteration by auto-configuring architectures and hyperparameters from the data. Comparative studies show that properly configured U-Nets can outperform more complex neural network architectures for segmentation tasks [26], supporting our emphasis on data quality over architectural novelty. In subsequent sections, we demonstrate that representative synthetic data is key to training a generalizable membrane segmentation model.

#### 2.1.1 Matching membrane density in experimental tomograms (V1–2)

Theoretically, PolNet can generate tomograms with an arbitrary density for each type of structure, like membranes. The definition of density here is equivalent to occupancy, the fraction of voxels occupied by the structure. Nevertheless, initially PolNet could not reproduce directly in practice the high occupancy of membranes present in some *in situ* tomograms of the validation set, described in Methods 4.1. The averaged occupancy estimated from the experimental validation set is 3.3 %, but PolNet computation times were unaffordable for densities greater than 2.5 %. In its original implementation [21], PolNet is based on trial-and-error and applies rigid transformations to membranes when it tries to insert a membrane. Therefore, its computational complexity scales with the tomogram size and the occupancy. To increase the membrane occupancy and reduce the computational complexity, we construct tomograms by composing a single tomogram of size 1000 × 1000 × 250 voxels with four tomograms of size 500 × 500 × 250 voxels. This approximates the 4 times binned tomograms from micrographs acquired at pixel sizes of 2-3 Å/px, typically used for segmentation. We achieve an occupancy of 3.5 % for membranes with this approach, which is equivalent to the occupancy found in the experimental validation set.

We trained model V1 on a synthetic training set at membrane occupancy 2.5 %, and model V2 at 3.5 %. Increasing membrane occupancy yields a higher midsurface Dice score (msDice), driven primarily by a precision gain (Fig. 2.B). We therefore retain the average occupancy of V2 for the subsequent cycles (V3-4). The synthetic training sets for V3 and V4 are detailed in the following sections. All performance scores are computed on the experimental validation set. The improvement of V2 is more pronounced on tomograms with high membrane occupancy; an example is shown in Fig. 3. We report msDice rather than standard Dice because it better reflects membrane topology and is less sensitive to membrane thickness; the details about its computation are provided in Methods 4.3.

**Fig. 3.**
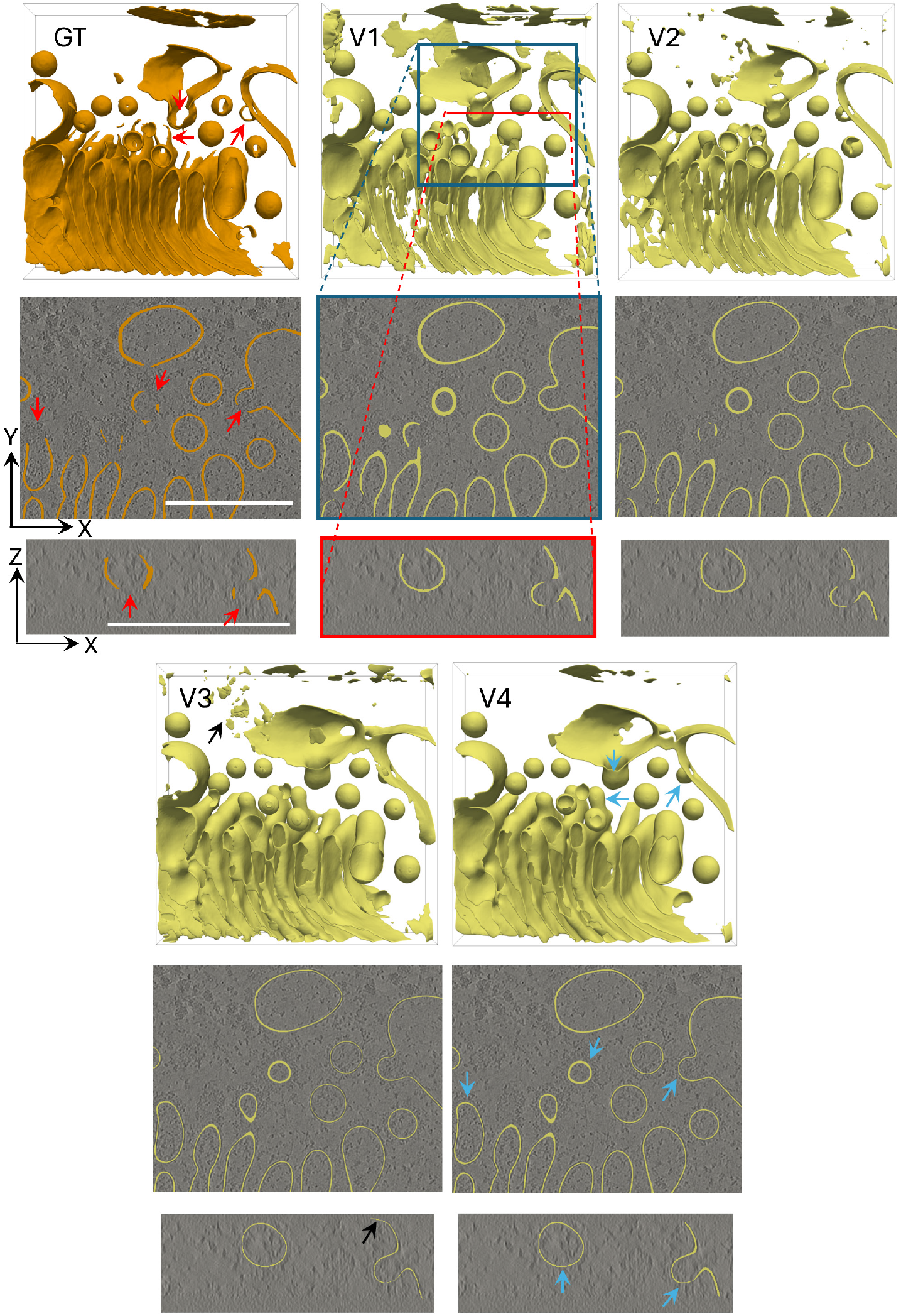
Visual comparison of the different TomoSegNet model versions (V1-V4) and ground truth (GT) of an experimental tomogram. Each result shows a 3D view of the full tomogram segmentation, along with two 2D slices of the tomogram region in the XY-plane (blue box) and the XZ-plane (red box). Red arrows point to examples of vanished membranes due to missing wedge, blue ones show how these membranes have been fully recovered in V4. Black arrows point to some oversegmentation examples of V3. Scale bars: 200 nm.

#### 2.1.2 Realistic membrane morphologies with curvatubes (V3)

PolNet initially simulated membranes using three simple geometries—spheres, ellipsoids, and tori—chosen for their mathematical tractability and their good coverage of local curvature combinations. Models V1-2, trained exclusively on these simple geometrical shapes, performed well on vesicles but were limited on organelles with more complex geometry and topology, such as the Golgi Apparatus, as shown in Fig. 3.

To address this limitation, we integrate the Curvatubes framework [27], which captures more complex and realistic membrane morphologies, into PolNet (more details in Sec. 2.4). We generate a new synthetic dataset containing solely membranes generated from Curvatubes for training V3. Relative to V2, V3 increases msRecall (0.65 → 0.73), but decreases msPrecision (0.93 → 0.79), yielding a similar msDice, see Fig. 2.B. The recall gain reflects improved segmentation of complex structures like the Golgi, illustrated in Fig. 3. The precision drop has two main causes: (1) V3 recovers membranes obscured by the missing wedge artifact that are absent from the expert annotations, and (2) V3 is trained on tomograms with a high concentration of intricate membranes, which induces a tendency for oversegmentation. The new method for generating membranes adds a higher diversity of shapes, thus enabling the segmentation model to recover more missing-wedge membranes. However, it also generates a higher density of membranes, specifically, in V3, we achieve an average occupancy of 8.9 %, far above most of the experimental tomograms. These observations motivated adjustments in V4 to recover precision without sacrificing recall.

#### 2.1.3 Diversity and distractors (V4)

Following the analysis of oversegmentation errors in V3, we broadened the training set distribution by combining the training sets of V3 and V2, yielding a mix of synthetic membranes (simple geometries and Curvatubes). We also included tomograms without membranes that contain only distractors (e.g., cytosolic proteins). Trained on this more diverse dataset, V4 substantially improves both recall (0.73 → 0.84) and precision (0.79 → 0.91) and shows more stable performance across the experimental validation tomograms (Fig. 2.B). Like V3, V4 can recover membranes that elude manual annotation, but it is less prone to segment non-membrane structures (Fig. 3).

Across V1–4, three levers—matching membrane occupancy of experimental tomograms, integrating the Curvatube algorithm to PolNet, and increasing training set diversity—yield consistent membrane segmentation improvement on the validation set. Notably, experimental tomograms were used only for error analysis: the model was trained exclusively on synthetic data, yet it performs well on experimental tomograms. Moreover, the visual comparison with reference segmentation, using ground truth, demonstrates the ability of TomoSegNet to recover membranes that vanished due to the missing wedge. The recovered membranes are not present in the ground truth. Consequently, a reduction in the msPrecision does not imply a worse prediction. In this case, it is simply a consequence of the impossibility of having a complete ground truth segmentation in cryo-ET.

### 2.2 Synthetic data are sufficient to train generalizable models

After summarizing iteration gains (V1-4) in Fig. 2, we compare the final model (V4) against state-of-the-art methods: MemBrain-Seg and TARDIS. Both are trained on diverse sets of experimental tomograms.

#### 2.2.1 Quantitative evaluation on the experimental validation set

We report quantitative performance on the experimental validation set (a subset of EMPIAR-11830) for which we have access to expert annotations. V4 was trained exclusively on synthetic data, although validation errors informed synthetic-data design. In contrast, MemBrain-Seg (1) was trained on a subset of EMPIAR-11830 and (2) was used to generate initial segmentations that the expert then curated into the reference annotations. This overlap limits the fairness of direct comparisons and helps explain MemBrain-Seg’s near-perfect scores.

On this validation set, TomoSegNet trained on V4 achieves competitive msDice, msPrecision, and msRecall relative to TARDIS and MemBrain-Seg (Fig. 4.A). All methods have comparable msPrecision. However, msRecall differs: 0.77 for TARDIS, 0.83 for TomoSegNet (V4), and 0.98 for MemBrain-Seg. Per-tomogram msRecall (Fig. 4.B) reveals two regimes: TomoSegNet (V4) performs strongly on vesicular/tubular membrane structures (tomograms 50, 167, 298, 649), approaching MemBrain-Seg, whereas performance drops on thylakoid-rich tomograms (closely stacked membrane sheets: 909, 929, 2162, 2917). MemBrain-Seg’s strength on thylakoids is expected, given their prevalence in its training data and its role in producing the initial annotations. V4 generalizes well to experimental tomograms, with the strongest performance on membrane morphologies that are adequately modeled and observed in the synthetic training distribution. In the next section, we examine experimental tomograms from different cell species and acquisition settings to further assess generalizability.

**Fig. 4.**
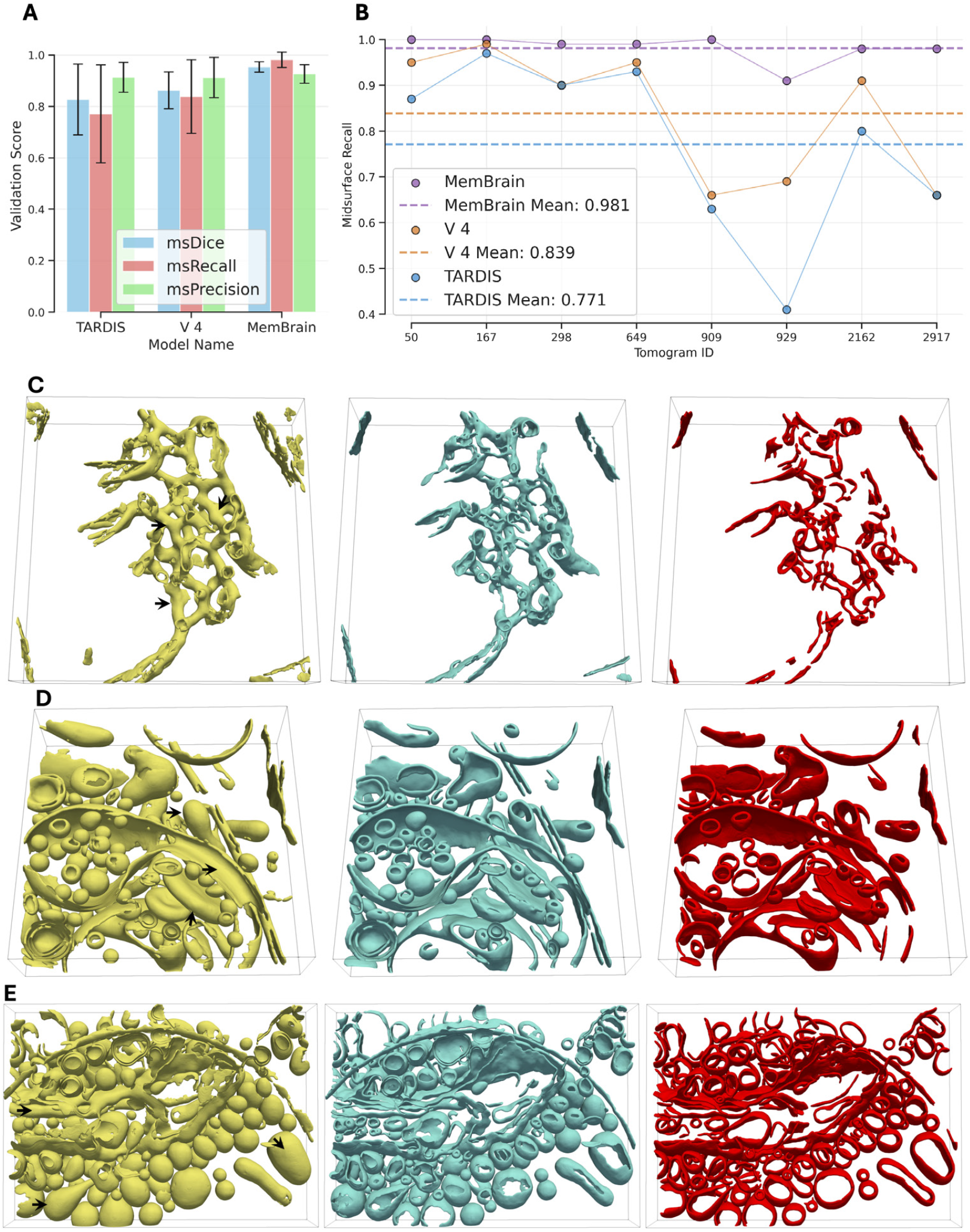
Comparing TomoSegNet (V4) to MemBrain-Seg and TARDIS. (A) msDice, msRecall, msPrecision for the experimental validation set, taken from EMPIAR-11830, which was used to train MemBrain-Seg. (B) Per-tomogram msRecall. (C-D) Qualitative evaluation on the experimental test set. TomoSegNet yellow, MemBrain-Seg blue, and TARDIS red. The arrows point to examples of missing wedge membranes recovered by TomoSegNet, and not reconstructed by the other methods. (C) EMD-12749. (D) EMD-50605. (E) EMD-15407.

#### 2.2.2 Qualitative evaluation on the experimental test set

Having established quantitative performance on the annotated validation set, we next assess generalizability qualitatively on a diverse experimental test set. To probe TomoSegNet’s robustness across cell species and acquisition settings, we selected publicly available tomograms, specially picked by MemBrain-Seg paper to showcase their model capabilities. These tomograms are suitable for the comparison because they were not included in the training data of MemBrain-Seg and were not examined or analyzed during the successive training iterations of TomoSegNet, ensuring that they remained truly unseen by both the model and the modelers. This separation between development and testing minimizes potential bias or information leakage from the validation set.

Reference annotations are unavailable for these tomograms; therefore our comparison is based on an expert’s visual assessment. For this qualitative evaluation we selected tomograms covering diverse membrane arrangements. Fig. 4.C shows a near-complete recovery of the intricate tubular network of pyrenoids. Fig. 4.E-D illustrate strong performance on vesicular membrane and more complex membranous organelles.

The visual comparison of the three methods revealed several key findings: (1) All models perform well on larger, well-defined membrane structures with comparable segmentation quality. (2) TomoSegNet consistently demonstrates superior performance in completing membrane structures affected by the missing wedge artifact, particularly in closing vesicles and predicting curved membrane closures that are not visible in the original data. TARDIS shows a poorer ability to compensate missing wedge. However, the evaluation also identified some limitations: (4) All models occasionally misclassify non-membrane structures as membranes, like the carbon hole, though TARDIS has a higher ability to suppress microtubules. (5) In some cases, TomoSegNet produces more false positives, showing a higher tendency to segment membrane-like small patches.

The performance of TomoSegNet is particularly notable given that it is trained solely on synthetic data generated by PolNet, suggesting that the physical models used to simulate the diverse and realistic membranes capture the complexities and challenges of real-world membrane segmentation tasks. These results demonstrate that our synthetic dataset is sufficient to train a model that generalizes well to real experimental data, even when the training data does not include any real tomograms.

### 2.3 Membranes recovered from missing-wedge regions

One motivation for training our model with synthetic data is to increase its ability to segment membranes hidden by the missing-wedge artifact, hereafter referred to as missing-wedge membranes. Synthetic data provides ground-truth annotations in this occluded region, a key advantage over experimental data, where these membranes are not visible and therefore cannot be annotated by experts. Our results from the previous section support this hypothesis, particularly when visually comparing the segmentation produced by our model with that from models mainly trained on experimental data.

#### 2.3.1 Quantitative evaluation of missing-wedge membranes

To enable further assessment of missing-wedge membrane recovery, we separate missing wedge membranes from the rest. To quantitatively assess TomoSegNet, we use unseen synthetic tomograms from a different distribution (large volume, low-density of membranes with a complex shape) and compute the evaluation metrics specifically for the missing-wedge membranes. In contrast, experimental tomograms only allow qualitative evaluation, since missing-wedge membranes lack ground-truth annotations.

As reported in Fig. 5, all models perform well on segmenting visible membranes: msDice of 0.96 for TomoSegNet, 0.94 for MemBrain-Seg, and 0.82 for TARDIS. We see a clear advantage for TomoSegNet when it comes to segmenting missing-wedge membranes: msDice of 0.88 for TomoSegNet, 0.30 for MemBrain-Seg, and 0.10 for TARDIS. MemBrain-Seg recovers parts of the missing-wedge membranes thanks to a specialized data augmentation technique, which applies a varying angular wedge in the Fourier domain to the gray-scale tomogram to increase its model robustness against the missing wedge. TomoSegNet recovers a good part of the missing-wedge membranes (msRecall of 0.85) with high precision (msPrecision of 0.92).

**Fig. 5.**
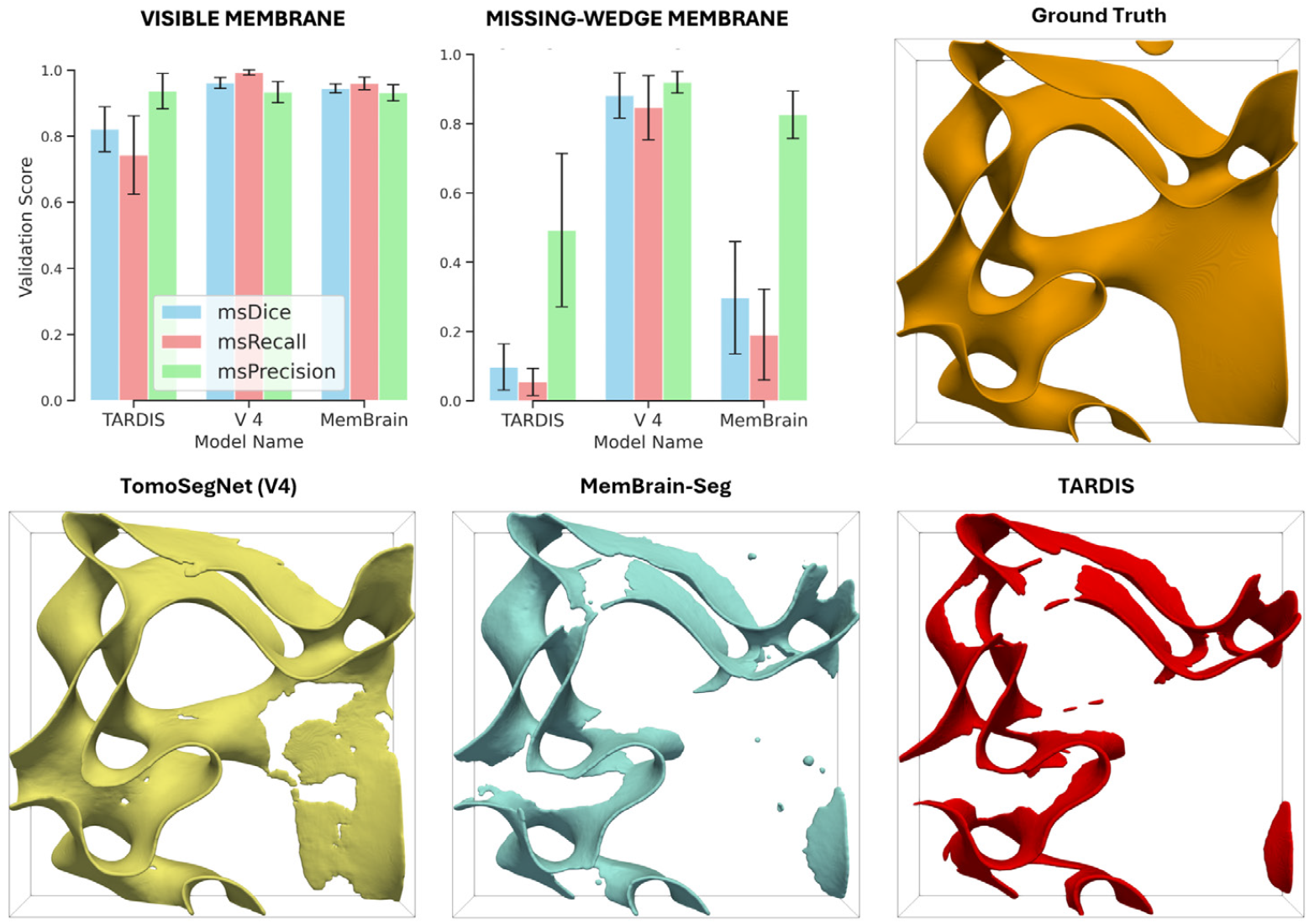
Recovering missing-wedge membranes, quantitative analysis. Comparing quantitatively and visually the segmentation performance of TomoSegNet (V4) with other methods. Non-missing wedge membranes are in orange for the ground truth, yellow for TomoSegNet, blue for MemBrain-Seg, and red for TARDIS, missing wedge membranes are in pink.

#### 2.3.2 Qualitative evaluation of missing-wedge membranes

In Fig. 6.A, we show 3D visualizations of the predictions from TomoSegNet and from MemBrain-Seg on experimental data. Interestingly, MemBrain-Seg prediction recovers some fractions of missing wedge membranes. Nevertheless, the amount of recovered membranes is considerably higher for TomoSegNet. In addition, tomogram slices along Z-axis (Fig. 6.C-L), the missing wedge direction, show how our model completes the missing membranes, in structures with a simple geometry like vesicles, as well as complex membranous organelles like the Golgi apparatus. Conversely, MemBrain-Seg cannot complete these membranes and, in many cases, tends to elongate membrane thickness along Z-axis, an artifact we suppose is produced by missing wedge.

**Fig. 6.**
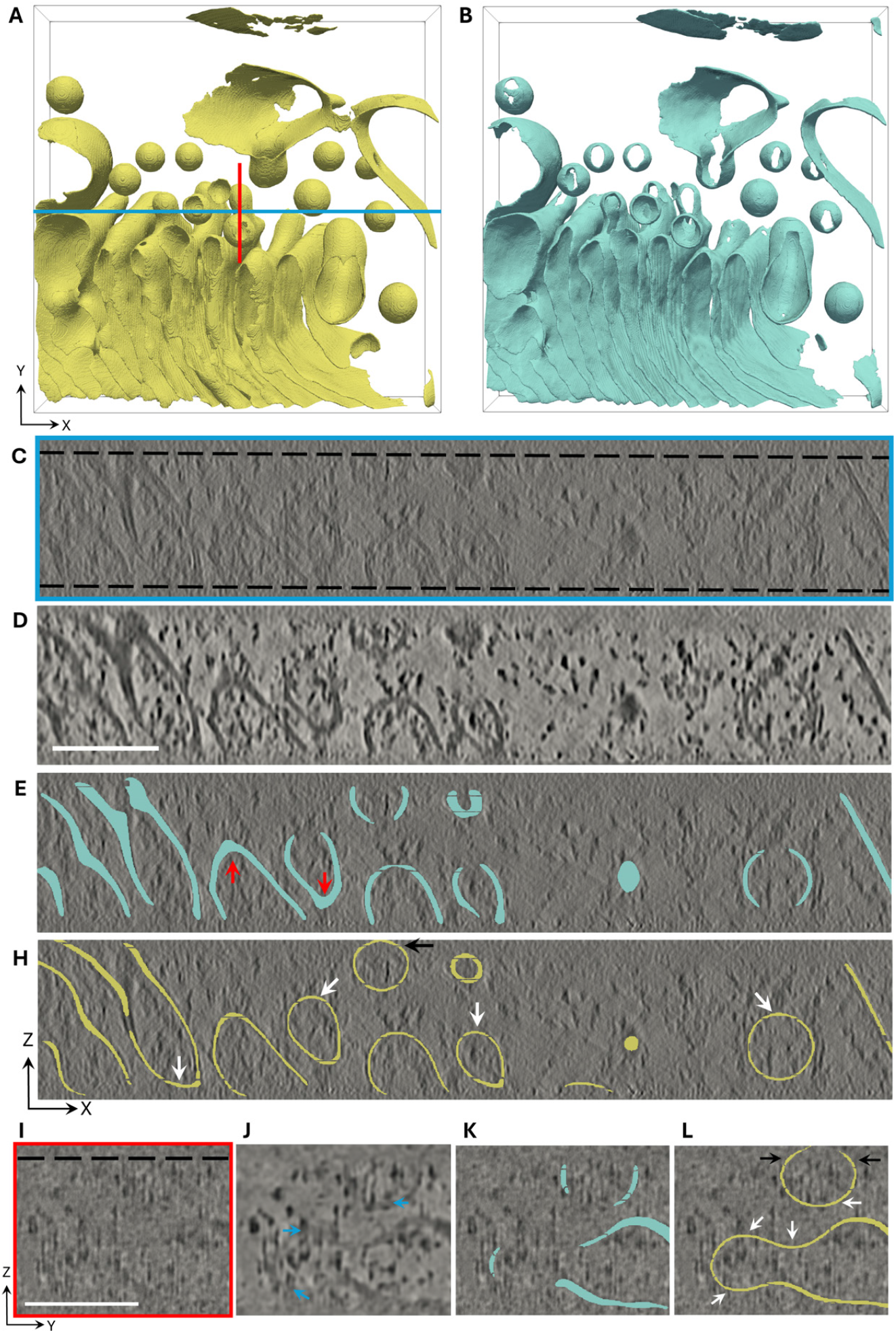
Recovering missing-wedge membranes, qualitative analysis. (A) TomoSegNet segmentation in 3D. (B) MemBrain-Seg segmentation of the same data used in A. (C) 2D XZ slice of the tomogram along the blue line in A. (D) IsoNet reconstruction of C. (E) MemBrain-Seg segmentation of C. (H) TomoSegNet segmentation of C. (I) 2D YZ slice of the tomogram along the red line in A. (J) IsoNet reconstruction of I. (K) MemBrain-Seg segmentation of I. (L) TomoSegNet segmentation of I. The dashed black lines enclose the lamella. White arrows point to examples of missing wedge membranes recovered, black arrows to examples of membrane predictions of the lamella, and blue ones to examples of membrane-bound proteins indicating the presence of membranes in missing wedge regions. Scale bars: 100 nm.

We hypothesize that TomoSegNet extrapolates the membranes in the missing wedge regions by analyzing the visible membranes and observing guiding features, like membrane-bound proteins. The membrane shape is predicted using the information learned during training from the biophysically informed simulated data. A crucial question is to determine whether the missing wedge membranes recovered by our model are plausible. It is impossible to validate the recovered membranes by visual comparison, as these membranes are not visible in the input real tomograms. Therefore, we speak about “plausible” membranes instead of “real” ones. For example, in Z-axis views of Fig. 6.H and L, we can see how our model predicts some membranes outside of the lamella. Obviously, in the imaged biological sample, these membranes do not exist because they are milled away during the sample preparation process. Nevertheless, the predicted membranes seem to provide a plausible approximation of the membranes in the sample before milling. We use two strategies to further assess the recovered missing-wedge membranes. First, we compare our predictions with those produced by IsoNet [28], a self-supervised deep learning model that restores the missing wedge directly in the input tomogram. Overall, IsoNet predictions are consistent with the missing-wedge membranes predicted by our model, although TomoSegNet recovers substantially more missing membranes, see Fig. 6.D, H, J and L. Second, we examine visible guiding features, such as membrane-bound proteins, to verify that they are embedded in the predicted membrane surfaces, see Fig. 6.J and L.

### 2.4 Diverse and complex synthetic membranes

Back in 1973, W. Helfrich [29] studied the elastic properties of lipid bilayers and proposed a geometric formulation of membrane bending energy in biological systems derived from physical principles. Just like stretching a rubber band requires more force than leaving it round, membranes pay an energetic cost when they bend. He theorized that the shape of closed lipid bilayer vesicles is governed by curvature elasticity (the resistance of a curved structure to bending), assuming the membranes behave as two-dimensional fluids that contain a constant volume. This led to a continuum theory of elasticity for membranes in which curvature emerges as the dominant contributor to the membrane’s free energy. We adapted this theory to generate synthetic membranes reproducing the high diversity of shapes present in real cellular membranes.

#### 2.4.1 Membrane generation

Helfrich modeled the membrane as a smooth, compact, and orientable surface, and postulated a specific bending energy functional. According to Helfrich, biological membranes are assumed to minimize this energy. The Curvatubes framework [27] generalizes the Helfrich energy into a class of curvature-based functionals of the form of a second-degree polynomial in the principal curvatures. In order to represent the surfaces in a discrete 3D grid, the energy functional is approximated using a phase-field formulation. A phase field is a smooth function used to represent boundaries between two phases. To numerically minimize the functional, the minimization problem is reformulated as a mass-preserving gradient flow, see Fig. 7.A. Therefore, the volume contained by the surface is approximately constant. This process is simulated by a GPU-accelerated algorithm that iteratively approximates an implicit solution for the problem.

**Fig. 7.**
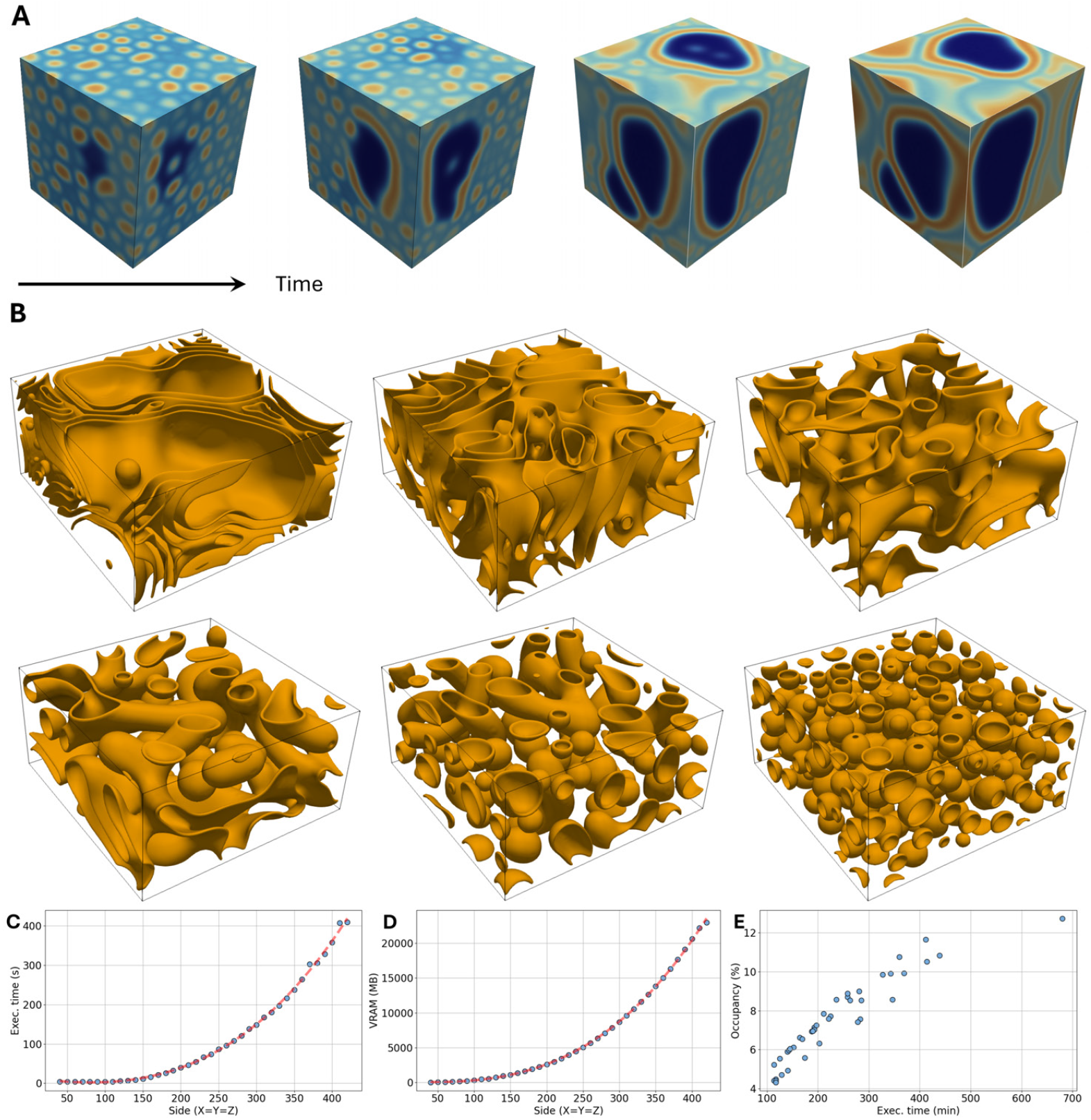
Generating membranes with complex geometries and topologies. (A) Convergence of the algorithm, an instance of the phase field over time. (B) Examples of different membrane shapes. (C) Run-time. (D) Spatial complexity. (E) Correlation between membrane occupancy and run-time. In C-E, blue dots represent experimental measures, and the dashed red lines represent the regressed polynomials; more details are given in Supplementary Information (Tabs. S1, S2, S3).

PolNet simulator was extended to include the algorithm of Curvatubes, which provides a final phase field per tomogram. Afterwards, the distance transform from the zero-thresholded phase field is computed. PolNet emulates the lipid bilayer inner structure of membranes using a double Gaussian profile along membrane normals. Finally, the distance information to the membrane surface for every voxel is used to generate the bilayer profile. A detailed description of the integration of Curvature in PolNet is available in Methods 4.4.

From the Helfrich functional, it is possible to intuitively derive global properties of the compact surfaces that minimize an energy functional. Based on these theoretical considerations, an exploratory parameter analysis was conducted to identify coefficient configurations that might generate membrane geometries and topologies far more complex and realistic than those in the original PolNet package, see Fig. 7.B. The details of this analysis are described in Methods 4.5. Despite not being exhaustive, this analysis provides a clear picture of how the parameters influence the nature of the generated surfaces.

#### 2.4.2 Computational performance analysis

One of the main limitations of the algorithm lies in its resource consumption, both in terms of time and memory. To measure this, the main routine for generating the membrane shapes was executed for a fixed number of iterations (1000) while varying only the side length of the cubic volume in which the surface is embedded. The results are shown in Fig. 7.C-D. Both cases follow a cubic trend concerning the side length of the volume; the correlation coefficient *R*^2^ is higher than 0.99 for both time and VRAM. This cubic growth behavior imposes a practical limitation on the size of tomograms that can be generated using the extended version of PolNet, especially considering that the GPU VRAM memory is typically limited to around 30 GB nowadays. However, this size limitation can be addressed by rescaling the phase field before generating the bilayer, thereby achieving run-times comparable to those of PolNet when simulating simple membranes or adding membrane-bound proteins for a tomogram at segmentation resolution around 10 Å per voxel.

Previous experiments show the relevance of simulating volumes with different membrane occupancies. Although the new approach for generating membranes cannot control membrane occupancy directly, the coefficient configurations proposed in Methods 4.5 obtain a high diversity of membrane occupancies. We found a direct correlation between the membrane occupancy and the running times, see Fig. 7.E.

## 3 Discussion

This work presents a data-driven approach that shows how current limitations for membrane segmentation in cryo-ET can be overcome by using machine learning and synthetic data. The advantages of using synthetic data are threefold: (1) there is no need for annotation of experimental data; (2) training a generalizable model capable of processing tomograms from different biological samples; and (1) recovering membrane areas vanishing due to the missing wedge.

Generating a diverse dataset is the key to training a generalizable and robust model, including complex geometries and topologies resembling the intricate membranous organelles of the cell. The model has to predict membranes relying on the partial information available as a result of the missing wedge. These predictions are plausible based on the fact that the model learned how to solve these situations from bio-physically grounded simulated data. Although we cannot ensure that these are real membranes, they seem to be good estimations considering the information available, that is, the visible membranes. Synthetic data can be used to quantitatively evaluate the power of our approach to recover missing wedge membranes. Nevertheless, a qualitative validation with real data is necessary. The careful observation of the recovered membranes, considering guiding features such as membrane-bound proteins, suggests that, in general, the predicted missing wedge membranes are plausible.

Besides diversity, reproducing the conditions of experimental tomograms is an important factor. Achieving a high occupancy of membranes helps to improve the results for processing real data, since tomograms enclosing membranous organelles may contain a high occupancy of membranes. Generating tomograms containing only a high occupancy of convoluted membranes improves the recall, but the predictions of a model trained only with these data are prone to oversegmentation. This behavior can be corrected through a diverse representation by adding tomograms with low occupancy, and even without membranes.

According to the paradigm proposed in this work, the challenge now is to transfer the knowledge accumulated by experimental sciences to machine learning algorithms. Here, the simulator is the instrument for transferring this knowledge, its precision and versatility ultimately determine the performance of the segmentation methods. In this line, our future efforts will focus on further improving the diversity and realism of the simulated membranes. Specifically, we want to improve the representation of the inner structure of the cellular membranes and include interactions between membranes.

## 4 Methods

### 4.1 Experimental validation set

A purpose of this work is to demonstrate that training membrane segmentation models solely from synthetic data has some advantages, including the recovery of membranes lost by distortions. However, we have also constructed a representative dataset of experimental, real, *in situ* tomograms to serve as references for parametrizing the synthetic simulator and evaluating the performance of the trained models.

For the sake of generalizing the results and quantitatively comparing with other publicly available methods, our validation set consists of eight tomograms of Chlamydomonas cells at 7.84 Å voxel size (EMPIAR 11830) that were expertly curated. These tomograms contain a diverse variety of membrane structures and arrangements, as shown in Fig. 3.

The expert annotation process employed a two-step approach: first, MemBrain-Seg was applied to generate initial membrane segmentations, as it was trained on tomograms from the same EMPIAR 11830 dataset. These segmentations were then manually refined and corrected by an expert to create the best possible annotations to be used as ground-truth. This setup allows us to evaluate whether TomoSegNet, trained exclusively on synthetic data, can match or exceed the performance of a state-of-the-art membrane segmentation tool that was trained on experimental data from the same source.

### 4.2 Synthetic datasets

This subsection describes the synthetic datasets prepared to train the different versions of the segmentation model.

#### 4.2.1 Dataset V1

The number of tomograms is 10 with a tomogram shape of 1000 × 1000 × 250 voxels, and a voxel size of 10 Å. The membrane occupancy is 2.5 %, and membranes were generated using basic shapes: spheres, ellipsoids, and toroids. We added instances of the cytosolic macromolecules, and the membranes were decorated with membrane-bound ones. The list of macromolecules included with their PDB codes is available in Tab. S4.

#### 4.2.2 Dataset V2

The number of tomograms is 20 with a tomogram shape of 500 × 500 × 250 voxels, and a voxel size of 10 Å. The membrane occupancy is 3.3 %, and membranes were generated using basic shapes: spheres, ellipsoids, and toroids. We added instances of the cytosolic macromolecules, and the membranes were decorated with membrane-bound ones. The list of macromolecules included with their PDB codes is available in Tab. S4.

#### 4.2.3 Dataset V3

The number of tomograms is 30 with a tomogram shape of 500 × 500 × 250 voxels, and a voxel size of 10 Å. The membrane occupancy is 8.9 %, and membranes were generated using the realistic approach described in Sec. 2.4. We added instances of the cytosolic macromolecules, and the membranes were decorated with membrane-bound ones. The list of macromolecules included with their PDB codes is available in Tab. S4.

#### 4.2.4 Dataset V4

The number of tomograms is 54 with a tomogram shape of 500 × 500 × 250 voxels, and a voxel size of 10 Å. The generation of membranes was divided into three sets. First, 20 tomograms with a membrane occupancy of 3.5 % and basic shapes. Second, 30 tomograms with realistic cellular membranes. Third, 4 tomograms without membranes. We added instances of the cytosolic macromolecules, and the membranes, when present, were decorated with membrane-bound ones. The list of macromolecules included with their PDB codes is available in Tab. S4.

### 4.3 Mid-surface Dice

We use the midsurface-Dice (msDice or *D*^*ms*^) instead of the standard Dice because it is more suited for membrane-like segmentations. Indeed, msDice focuses on the topological correctness of the predicted segmentation, ignoring that membrane thickness can be misleading when using pixel-wise accuracy. To compute the msDice, we need to first extract the midsurface of the predicted volume and the midsurface of the ground truth volume. The midsurface corresponds to a 3D skeleton of the segmented membrane, a thinned version of the volume that should be centered relative to the volume’s boundary.

Let us define *V*_*GT*_ as the ground truth volume and 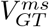 as the midsurface of the ground truth volume. Similarly, *V*_*P*_ is the predicted volume and 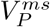 its corresponding midsurface. We compute the midsurface recall, msRecall or *R*^*ms*^, and the midsurface precision, msPrecision or *P* ^*ms*^, as follows:

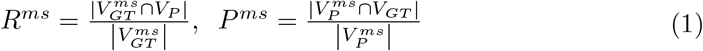

Finally, the msDice corresponds to the harmonic mean of the midsurface recall and the midsurface precision:

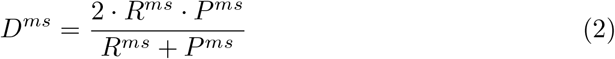

This metric was originally introduced as “surface dice” by Lamm et al. [17] for membrane segmentation, adapting the centerline dice metric [30] that was designed for tubular-like structure segmentation in 2D images. However, we identify a potential terminological ambiguity: the term “surface dice” has been previously established in medical imaging literature [31] as a metric that evaluates the overlap between two boundary surfaces within a specified tolerance distance. This differs fundamentally from the membrane-specific metric proposed by Lamm et al., which operates on the midsurfaces of volumetric membrane structures rather than their boundaries. To avoid confusion and clearly distinguish between these two distinct metrics, we adopt the more descriptive term “midsurface dice” throughout our work.

### 4.4 Integration of Curvatubes into PolNet

In contrast to the parametric surface generation method, Curvatubes utilizes an implicit representation of surfaces. We therefore had to adapt the existing software of PolNet https://github.com/anmartinezs/polnet.git to produce membranes with the same characteristics.

We began by creating a package to contain the Curvatubes `cvtub` module (https://github.com/annasongmaths/curvatubes.git). This module comprises:

- `curvdiags.py`: procedures to compute and plot the curvature diagram of a surface 𝒮 = {*u* = 0} defined by a phase-field *u*.
- `energy.py`: includes the process to compute the diffuse curvature energy *F*_*ϵ*_(*u*). We used GPU-accelerated PyTorch operations for efficient computation.
- `filters.py`: contains the methods to apply Gaussian filters and differential operators such as gradient, Hessian, or divergence.
- `generator.py`: defines the main wrapper function for generating the surfaces.
- `utils.py`: auxiliary functions used throughout the rest of the framework. It includes I/O management, visualization tools, and initialization routines.

The `lrandom.py` file, which defines a parameter generator for each of the membrane classes, was extended to provide the necessary parameters for this type of surface. The role of these parameters is explained in Methods 4.5, and they are:

- `mass_rg`: mass of the volume 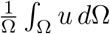.
- `a11_rg`: the range for the coefficient *a*_1,1_ in the energy functional.
- `a02_rg`: the range for the coefficient *a*_0,2_ in the energy functional expression.
- `b_rg`: the range for the coefficients *a*_1,0_ and *a*_0,1_ in the energy functional expression.
- `c_rg`: the range for the coefficient *a*_0,0_ in the energy functional expression.

In this way, we establish intervals where the parameters are uniformly selected. Additionally, we allow for their manual specification, which is recommended when searching for specific properties.

Finally, we have added a new membrane class, `MbCurvatubes`, within the `membrane.py` module. This class inherits from the pre-existing `MB` class. The procedure for generating the membrane begins by invoking the `_generate_shape` method from Curvatubes to create a scalar field *u*. Then, the intermediate volume `ct den` is defined as the boundary of the region where *u* ≤ 0. This is computed using the `binary_dilation` function from SciPy’s `ndimage` module. The binary mask `ct_den` represents the membrane’s skeleton. Afterward, we use the distance transform, also from `scipy.ndimage`, to encode the voxel-wise Euclidean distance to `ct_den` into `ct_dist`. The membrane mask is then extracted using the distance threshold defined by the membrane thickness (`mb_mask` = `ct_dist` < `thickness`). Lastly, the lipid bilayer included in the sample is obtained by subtracting the skeleton from the mask and applying a subsequent Gaussian filter to smooth the membrane profile.

#### Algorithm 1

Lipid bilayer generation from curvatubes output

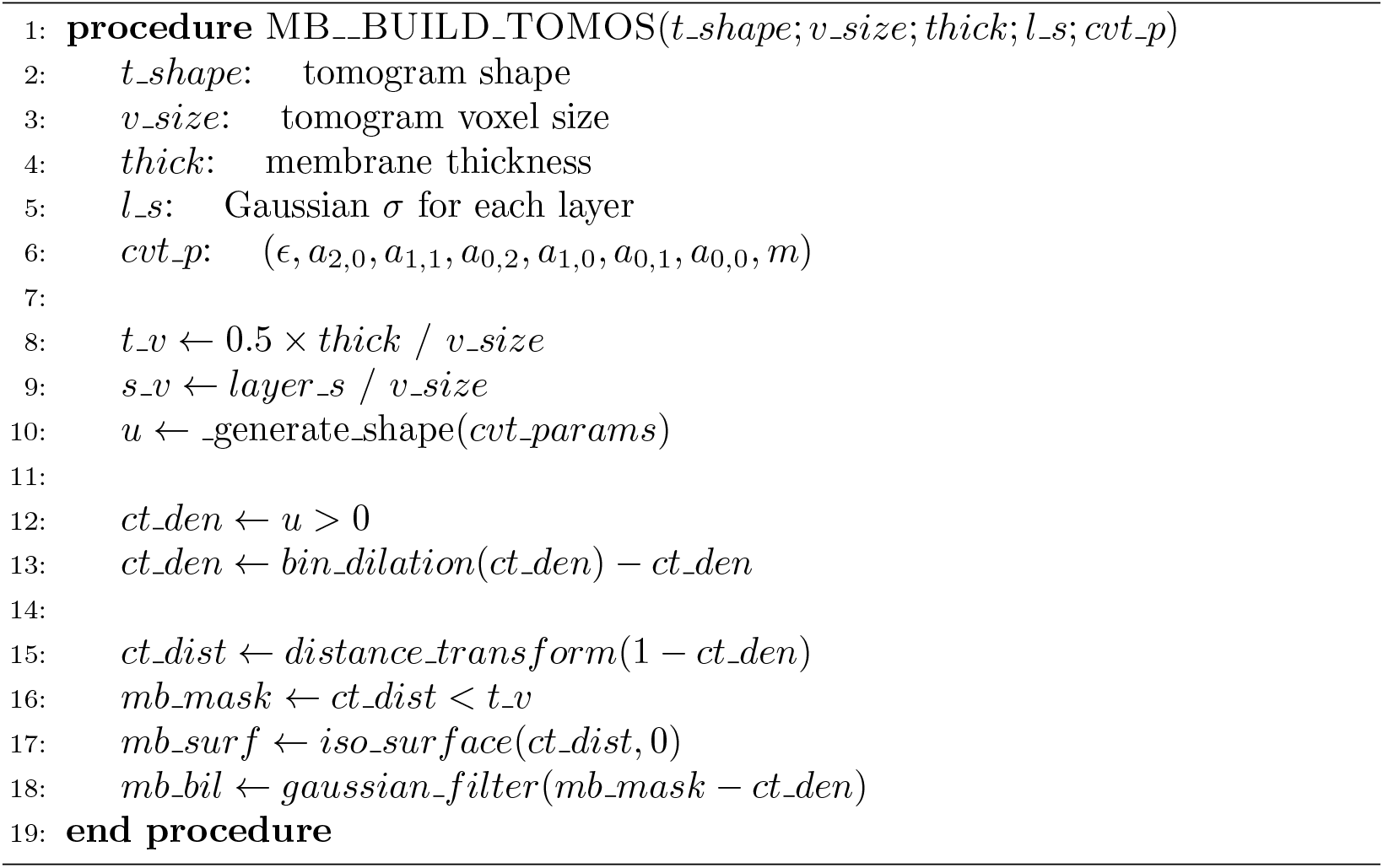

### 4.5 Parameter selection for Curvatubes

The generation of surfaces using Curvatubes essentially depends on seven parameters: the six coefficients of the polynomial *p*(*κ*_1_, *κ*_2_) and the initial mass parameter *m*_0_. We utilize the Willmore energy functional [32] as a baseline for our analysis. This corresponds to the parameter set (*a*_2,0_, *a*_1,1_, *a*_0,2_, *a*_1,0_, *a*_0,1_, *a*_0,0_) = (1, 1, 1, 0, 0, 0).

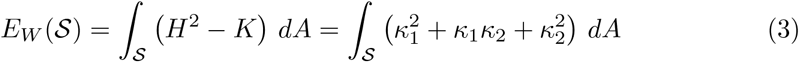

Note that we define the mean curvature as the sum of principal curvatures *H* = *κ*_1_ + *κ*_2_ rather than the average to maintain consistency with the definition used in the Curvatubes source code and reference article.

If we suppose that biological membranes are closed surfaces, we can use the Gauss-Bonnet Theorem to infer some properties that result from varying *λ* = *a*_1,1_

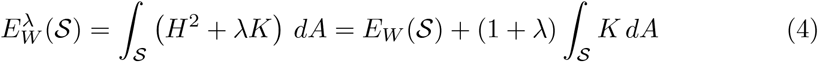

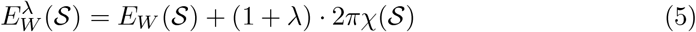

Here, *χ*(𝒮) denotes the Euler characteristic of the surface. Assuming the surface is orientable, this can be expressed as a function of its genus *g* (the number of “holes”) by *χ*(𝒮) = 2 −2*g*. Thus, the term (1+*λ*) · 2*π* · (2 −2*g*) modifies the energy output based on topology. Specifically, since the genus *g* contributes negatively to the Euler characteristic, a large positive value for (1 + *λ*) lowers the energy cost of high-genus surfaces. Consequently, increasing *λ* favors the formation of holes during the minimization process (sponges, tubular surfaces…), while values of *λ* such that (1 + *λ*) < 0 tend to produce topologically simple surfaces. In the context of Helfrich’s theory for elasticity, this coefficient corresponds to the Gaussian bending modulus of the membrane.

The linear coefficients *a*_1,0_ and *a*_0,1_ control the scale of the features. Following the analogy to the Helfrich model, introducing a target mean curvature *H*_0_ into the Willmore functional, the first term becomes

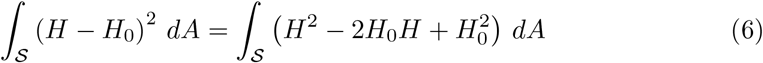

Note that the integral is zero if and only if *H* ≡ *H*_0_, so one part of the minimization process aims to globally approximate 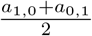. Setting these parameters to non-zero values encourages the formation of cylindrical or spherical structures with a preferred radius *R* roughly inversely proportional to the magnitude of the linear coefficients, i.e., 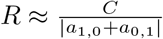.

The remaining quadratic terms *a*_2,0_ and *a*_0,2_ act as asymmetry factors. Nevertheless, as Helfrich suggested, to ensure physical plausibility for isotropic surfaces, the energy functional should be invariant under the exchange of principal curvatures. This theoretically imposes the constraints *a*_2,0_ = *a*_0,2_ and *a*_1,0_ = *a*_0,1_. The constant term *a*_0,0_ acts as the surface tension parameter, as the integral ∫_𝒮_*a*_0,0_ *dA* is proportional to the total surface area, penalizing high occupancy membranes.

Finally, the parameter *m*_0_ does not affect the geometric or topological properties of the surface per se, but rather the global volume constraint. In the context of the implicit representation of the surface in Curvatubes, *m*_0_ ∈ [−1, 1] represents the average value of the phase field.

Following this theoretical analysis, we experimented with the software generating 1250 different surfaces combining different values of these parameters:

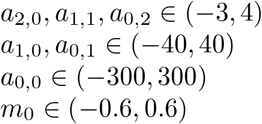

Then, we selected thirty of the resulting surfaces to incorporate into the Polnet-generated dataset.

## Acknowledgements

A.M-S. is supported by grants RYC2021-032626-I and CNS2023-144921 funded by MICIU/AEI/10.13039/501100011033 and the European Union by NextGenerationEU/PRTR, and the grant PID2023-151075OA-I00 funded by MICIU/AEI/10.13039/501100011033 and FEDER, UE. A.M-S. is also supported by grant FS/10.13039/100007801(22686/PI/24) funded by Fundaciôn Séneca -Agencia de Ciencia y Tecnología de la Regiôn de Murcia (Spain) and the Universidad de Murcia through its program AttractRyC 2023.

## Supplementary Information

**Table S1.** Execution metrics after 1000 iterations of Curvatubes algorithm.

| Side length ($X = Y = Z$) | Time after 1000 iterations (s) | Allocated VRAM (MB) |
| --- | --- | --- |
| 40 | 3.79 | 22 |
| 50 | 3.71 | 58 |
| 60 | 3.75 | 90 |
| 70 | 3.90 | 122 |
| 80 | 3.73 | 182 |
| 90 | 3.77 | 242 |
| 100 | 3.71 | 344 |
| 110 | 4.35 | 484 |
| 120 | 5.50 | 564 |
| 130 | 6.97 | 768 |
| 140 | 8.85 | 934 |
| 150 | 11.80 | 1138 |
| 160 | 15.57 | 1356 |
| 170 | 21.15 | 1642 |
| 180 | 26.22 | 1942 |
| 190 | 32.72 | 2278 |
| 200 | 39.69 | 2620 |
| 210 | 46.61 | 3006 |
| 220 | 54.78 | 3460 |
| 230 | 66.49 | 3964 |
| 240 | 74.25 | 4508 |
| 250 | 86.42 | 5060 |
| 260 | 95.92 | 5694 |
| 270 | 107.93 | 6372 |
| 280 | 120.52 | 7082 |
| 290 | 137.78 | 7884 |
| 300 | 147.92 | 8714 |
| 310 | 167.86 | 9600 |
| 320 | 180.38 | 10568 |
| 330 | 196.80 | 11586 |
| 340 | 215.93 | 12634 |
| 350 | 237.54 | 13812 |
| 360 | 264.32 | 14994 |
| 370 | 302.72 | 16302 |
| 380 | 304.84 | 17646 |
| 390 | 328.12 | 19106 |
| 400 | 357.08 | 20618 |
| 410 | 406.87 | 22176 |
| 420 | 408.89 | 22962 |

**Table S2.** Cubic fit coefficients for metrics in Tab. S1.

| *Metric* | $x^3$ | $x^2$ | $x^1$ | $x^0$ |
| --- | --- | --- | --- | --- |
| Exec. time | 4.49938435e-06 | 1.00435971e-03 | -2.52082076e-01 | 1.36769617e+01 |
| VRAM | 2.82622259e-04 | 2.16657459e-02 | -3.26758330e+00 | 1.53850877e+02 |

**Table S3.** Polnet execution time for 41 tomograms. Curvatubes algorithm was iterated 6000 times.

| Tomo | Occupancy(%) | Time(min) | Protein Insertion Time (%) | Pns | Pms |
| --- | --- | --- | --- | --- | --- |
| 0 | 8.56 | 346 | 9.25 | 9631 | 5461 |
| 1 | 6.61 | 164 | 19.51 | 9858 | 3080 |
| 2 | 4.48 | 117 | 27.35 | 10100 | 2922 |
| 3 | 7.57 | 283 | 11.31 | 9754 | 4899 |
| 4 | 7.13 | 193 | 16.58 | 9788 | 3374 |
| 5 | 9.93 | 369 | 8.67 | 9478 | 4715 |
| 6 | 5.89 | 140 | 22.86 | 9942 | 2801 |
| 7 | 4.4 | 114 | 28.07 | 10108 | 2886 |
| 8 | 8.53 | 285 | 11.23 | 9640 | 4068 |
| 9 | 7.71 | 225 | 14.22 | 9734 | 3658 |
| 10 | 10.84 | 439 | 7.29 | 9301 | 4982 |
| 11 | 6.93 | 188 | 17.02 | 9823 | 3297 |
| 12 | 8.99 | 281 | 11.39 | 9587 | 4229 |
| 13 | 10.52 | 414 | 7.73 | 9400 | 5004 |
| 14 | 8.7 | 258 | 12.4 | 9624 | 4030 |
| 15 | 8.53 | 263 | 12.17 | 9638 | 4058 |
| 16 | 4.41 | 117 | 27.35 | 10106 | 2919 |
| 17 | 10.76 | 360 | 8.89 | 9387 | 4957 |
| 18 | 5.53 | 125 | 25.6 | 9981 | 2634 |
| 19 | 8.88 | 258 | 12.4 | 9601 | 4087 |
| 20 | 7.58 | 221 | 14.48 | 9745 | 3611 |
| 21 | 5.23 | 114 | 28.07 | 10014 | 2490 |
| 22 | 4.92 | 141 | 22.7 | 10048 | 3201 |
| 23 | 7.85 | 211 | 15.17 | 9718 | 3607 |
| 24 | 4.71 | 129 | 24.81 | 10075 | 3077 |
| 25 | 7.25 | 197 | 16.24 | 9786 | 3456 |
| 26 | 5.58 | 174 | 18.39 | 9946 | 3784 |
| 27 | 5.96 | 143 | 22.38 | 9931 | 2982 |
| 28 | 12.74 | 680 | 4.71 | 9157 | 5898 |
| 29 | 7.03 | 193 | 16.58 | 9810 | 3344 |
| 30 | 6.32 | 203 | 15.76 | 9893 | 4093 |
| 31 | 9.86 | 327 | 9.79 | 94492 | 4557 |
| 32 | 11.65 | 412 | 7.77 | 9066 | 5351 |
| 33 | 6.55 | 169 | 18.93 | 9865 | 3124 |
| 34 | 6.11 | 152 | 21.05 | 9916 | 2890 |
| 35 | 7.42 | 278 | 11.51 | 9768 | 4876 |
| 36 | 6.95 | 190 | 16.84 | 9824 | 3305 |
| 37 | 9.91 | 343 | 9.33 | 9485 | 4649 |
| 38 | 6.04 | 145 | 22.07 | 9928 | 2888 |
| 39 | 4.31 | 117 | 27.35 | 10119 | 2830 |
| 40 | 8.56 | 236 | 13.56 | 9638 | 3940 |

**Table S4.**
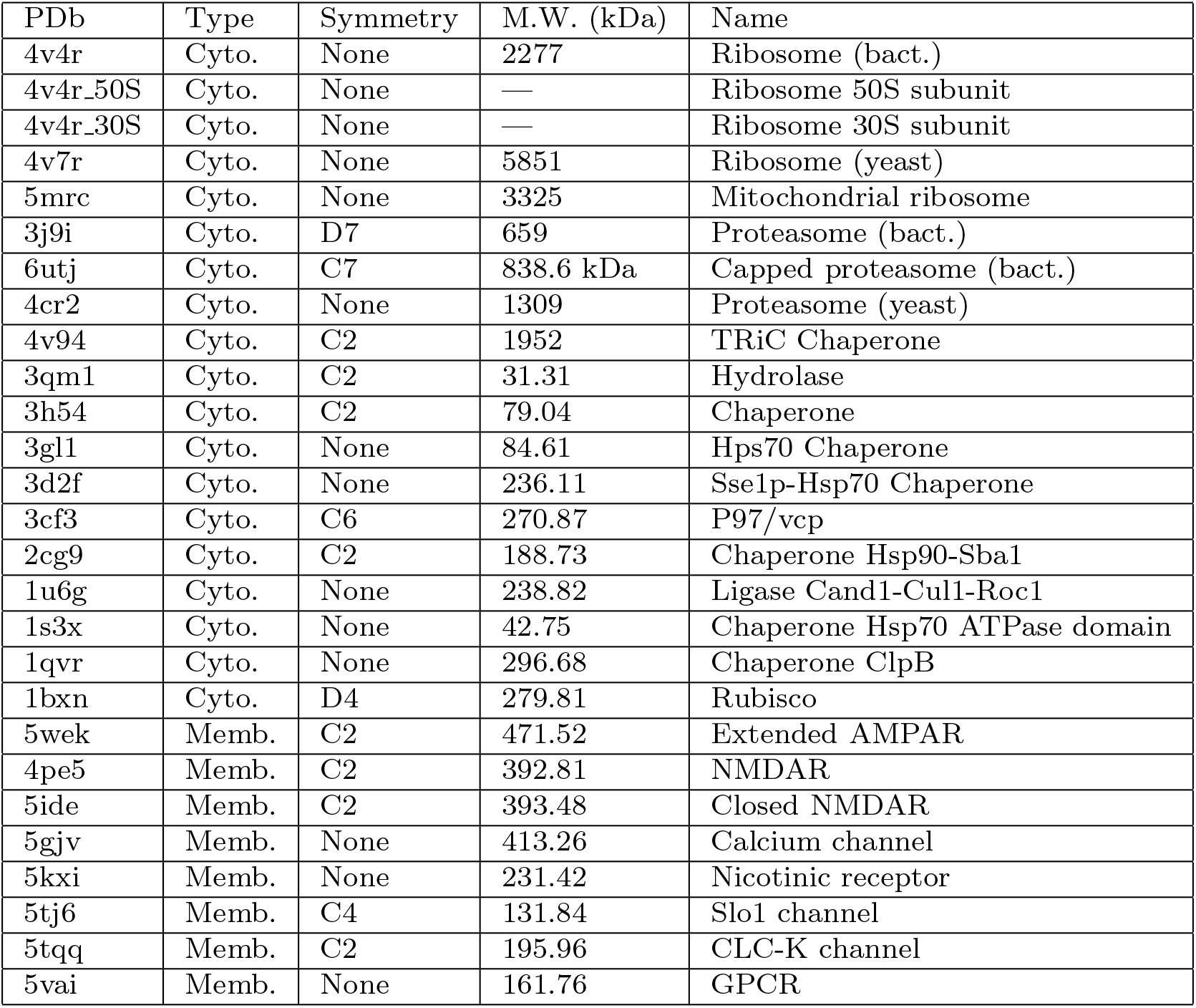
List of macromolecules included in simulations.

